# A Novel Genomic Rearrangement in the *Amaranthus palmeri* Extrachromosomal Circular DNA Provides Dual Herbicide Resistance to Glyphosate and Glufosinate

**DOI:** 10.1101/2024.10.08.617243

**Authors:** Pâmela Carvalho-Moore, Ednaldo A. Borgato, Luan Cutti, Aimone Porri, Ingo Meiners, Jens Lerchl, Jason K. Norsworthy, Eric L. Patterson

**Affiliations:** Crop, Soil and Environmental Science Department, University of Arkansas, Fayetteville, AR 72703, USA; Department of Plant, Soil, and Microbial Science, Michigan State University, East Lansing, Michigan 48823, USA; BASF SE, Ludwigshafen 67063, Germany; BASF Corporation, Research Triangle Park, NC 27709, USA

## Abstract

Amplification of chloroplastic glutamine synthetase (*GS2*) has been characterized as one of the resistance mechanisms in glufosinate-resistant *Amaranthus palmeri* accession (MSR2). Previously, the overamplification of the glyphosate-resistance gene, 5-enolpyruvylshikimate-3-phosphate (*EPSPS*), in *A. palmeri* was determined to be driven by an extrachromosomal circular DNA (eccDNA). Here, a novel eccDNA is described that carries both glyphosate and glufosinate-ammonium target site due to co-duplication of their chromosomic native region, conferring resistance. Besides *EPSPS*, the novel replicon has a region replaced by a fragment carrying the *GS2* isoforms (*GS2.1* and *GS2.2*) and other genes. The co-existence of eccDNA carrying only *EPSPS* was confirmed in MSR2 samples harboring dual targeting eccDNA. The genomic structure of *GS2* and *EPSPS* amplification was also assessed in a different glufosinate-resistant *A. palmeri* accession (MSR1) along with MSR2. The accessions showed distinct *GS2.1* and *GS2.2* amplification patterns suggesting the existence of diverse replicons that were not assembled here. The *EPSPS* was amplified in both accessions, and a correlation was observed with the *GS2* isoforms in MSR2, further supporting the co-existence of these genes in the same replicon. These findings shed light on the complexity of eccDNA formation in plant systems, with the collection and accumulation of extra pieces of DNA.

## Introduction

Numerous factors influence the speed of evolution, chief among these factors is sufficient genetic diversity within a population on which selection can act. Herbicide resistance evolution is a near-perfect example of evolution in action, and the speed at which it occurs can be surprising. Weeds have evolved several mechanisms to survive herbicides; single nucleotide polymorphisms (SNPs) in the herbicide target-site gene and the increased capacity of metabolizing the herbicide are among the most common (Powles and Yu 2010). However, several other mechanisms appear less frequently, such as genomic rearrangements that result in the over-expression of herbicide target proteins so-called copy number variation (CNV) (Gaines et al. 2010). To date, this mechanism is primarily found in glyphosate-resistant populations of both monocot and eudicot weeds.

The enzyme 5-enolpyruvylshikimate-3-phosphate synthase is the target site of the herbicide glyphosate (EPSPS; Steinrücken and Amrhein 1980). Increased gene copies of *EPSPS* result in increased *EPSPS*-mRNA and proportionally increased *EPSPS*-protein. It takes a greater amount of applied glyphosate to inhibit the increased EPSPS-protein pool; therefore, with only a few additional gene copies, weedy plants can resist typical field doses of glyphosate. Several species have evolved *EPSPS*-CNVs, each through unique structural rearrangements. Kochia (*Bassia scoparia*) has the *EPSPS* tandemly amplified at a single locus (Jugulam et al. 2014; Patterson et al. 2019); goosegrass (*Eleusine indica*) has a tandem *EPSPS* amplification in the subtelomere region of one of its chromosomes (Zhang et al. 2023); waterhemp (*Amaranthus tuberculatus*) has the *EPSPS* extra copies also in tandem duplication close to its native locus, near the centromere, but also extra copies on an extra chromosome (Dillon et al. 2017). However, the first CNV identified in herbicide-resistant weeds was reported in Palmer amaranth (*Amaranthus palmeri*; Gaines et al. 2010), and the most recent in Italian ryegrass (*Lolium perenne* ssp. *multiflorum*; Koo et al. 2023), showing the *EPSPS* being duplicated through a genome-independent, autonomously replicating, circular piece of DNA called extrachromosomal circular DNA (eccDNA) (Koo et al. 2018b). Thus far, the nature and origin of these unique structural rearrangements are not fully understood, with some hypotheses around transposable elements activity and unequal crossing over (Hall et al. 2023; Jugulam et al. 2014); however, now that researchers are investigating these structures more deeply, they seem to be quite common (Fu et al. 2023).

The first report of eccDNAs was from 1965 (Hotta and Bassel 1965). The functions and phenotypic consequences of eccDNAs are still being unveiled but are much better understood in humans than in plants, where many functions have been characterized (Zuo et al. 2022). For instance, an eccDNA can cause drug resistance in cancer cells as it carries both oncogenes and/or chemotherapy-resistance genes (Turner et al. 2017). In yeast, eccDNAs have been shown to carry the copper resistance gene *CUP1* under elevated environmental copper (Hull et al. 2019). Interestingly, there is a near-direct parallel to the *A. palmeri* eccDNA that confers herbicide resistance due to *EPSPS* amplification (Koo et al. 2018a) or the eccDNA in blackgrass (*Alopecurus myosuroides*) that has the herbicide metabolization gene *GSTF1* (Fu et al. 2023). EccDNA formation remains not fully comprehended. It has been hypothesized that stress might generate fragments of DNA which are consequently recognized and repaired by DNA damage-repairing mechanisms to form eccDNAs (Zuo et al. 2022) or that they are formed from the circularization of extrachromosomal linear DNA of active transposable elements (Møller et al. 2016), or even as random DNA rearrangement processes (Zhuang et al. 2024). Considering the first hypothesis, eccDNAs may rapidly emerge as a defense or resistance mechanism under physiological stress in eukaryotes, an attractive explanation for the rapid rate at which cancer drug and herbicide resistance can be accumulated (Luo et al. 2023; Molin et al. 2020b; Zhuang et al. 2024).

*Amaranthus palmeri* has become one of the most problematic weeds in row crop fields worldwide, causing yield losses of up to 91% in row crop production systems (Massinga et al. 2001). One of the reasons for its meteoric rise in importance is its ability to evolve resistance to herbicides, which shrinks the already limited postemergence options. To date, it has evolved resistance to almost all critical herbicides, such as glyphosate, 2,4-D, 4-hydroxyphenylpyruvate dioxygenase-, protoporphyrinogen oxidase-, and acetolactate synthase (ALS)-inhibitors, usually through independent mechanisms (Gaines et al. 2010; Hwang et al. 2023; Küpper et al. 2017; Varanasi et al. 2018). Rampant glyphosate resistance has led growers to lean more heavily on alternative chemistries that target multiple herbicide sites of action (Norsworthy et al. 2012). With the commercialization of glufosinate-resistant cotton (*Gossypium hirsutum*) and soybean (*Glycine max*), glufosinate-ammonium became foundational in chemical control programs targeting *A. palmeri* accessions carrying resistance to other herbicides (Norsworthy et al. 2016). This non-selective herbicide inhibits the enzyme glutamine synthetase (*GS*), which has two isoforms in plants, *GS1* in the cytosol and *GS2* in the chloroplast (Bayer et al. 1972; McNally and Stewart 1983). Glutamine synthetase catalyzes the condensation of glutamate and ammonia to form glutamine (Hoerlein 1994). The *GS* enzyme inhibition leads to ammonia buildup and reactive oxygen species overproduction, which subsequently leads to lipid peroxidation and plant death (Takano et al. 2020). As with other herbicides, recurrent glufosinate-ammonium application has likely led to resistance evolution in *A. palmeri* (Carvalho-Moore et al. 2022; Priess et al. 2022; Noguera et al. 2022). The mechanism of glufosinate-ammonium resistance in *A. palmeri* is still being described, and target- and non-target-site resistance mechanisms have been described among resistant weeds (Brunharo et al. 2019; Carvalho-Moore et al. 2022; Noguera et al. 2022; Zhang et al. 2022). Target-site mechanisms conferring glufosinate-ammonium resistance have been reported for Ser59Gly, in *GS1* in *E. indica* (Zhang et al. 2022) and *GS2* amplification in *A. palmeri* (Carvalho-Moore et al. 2022; Noguera et al. 2022).

The genomic structural rearrangement driving the *GS2* amplification is unknown in glufosinate-ammonium-resistant accessions. One of the hypotheses is that gene amplification of *GS2* is located in an eccDNA molecule, as reported for glyphosate resistance (Koo et al. 2018b). Therefore, the objective of the current work was to characterize the genomic rearrangement involved in *GS2* amplification in an *A. palmeri* accession resistant to both glufosinate-ammonium and glyphosate. Moreover, this study investigated the relationship between *GS2* CNV and eccDNA-driven amplification of the glyphosate target gene *EPSPS*.

## Results

### Identification of new eccDNA rearrangement *in A. palmeri*

In this study, it is reported a novel *A. palmeri* eccDNA that confers resistance to both glyphosate and glufosinate-ammonium by co-duplication of each target site in the same replicon. The glufosinate-ammonium-resistant accession MSR2 was previously characterized for having increased gene copy number and transcription of the glufosinate target *GS2* (Carvalho-Moore et al. 2022). Additionally, it has been confirmed that the same accession is also resistant to glyphosate (Supplemental Figure S1). To uncover the specific structural variant responsible for *GS2* amplification, a new eccDNA from shotgun, PacBio HiFi reads that is 426,133 bp long, 26,698 bp longer than the previous eccDNA identified in *A. palmeri* (Molin et al. 2020a) was assembled. The two eccDNAs show high levels of similarity and synteny (Figure 1); however, clearly, a region in the original eccDNA has been swapped for the locus that contains the tandemly duplicated *GS2.1* and *GS2.2* genes in the genome (Figure 2c). The fragment from 84,958 to 106,910 bp (21,952 bp total) of the previously known *A. palmeri* eccDNA was removed in the new assembly with a new 53,159 bp long sequence inserted in its place (Figure 2a). Most of the new sequence has synteny with chromosome 15 of the *A. palmeri* genome; however, a 5,476 bp long sequence at the beginning of the insertion and a 1,568 bp in the middle are unknown and do not have similarity with any sequence from the *A. palmeri* reference genome assembly. The inserted sequence before the 1,568 unknown fragment, which contains *GS2.1*, *GS2.2*, has 76.5% similarity with its native region in chromosome 15, while the sequence after has 87% similarity. This sequence replacement event in the new eccDNA did not affect the *EPSPS* gene locus, which also appeared co-duplicated in the same replicon ∼24,600 bp downstream of the chromosome 15 insertion (Figure 2b). The *EPSPS* from eccDNA shows synteny to its native location in chromosome 7 and was inserted with its 5,572 bp upstream and 1,124 bp downstream native region (Figure 2a). In addition, from the same individual in which the new eccDNA was discovered, it also had reads supporting the presence of the original replicon (with only *EPSPS*), which implies both the original and new eccDNAs co-exist and independently replicate (Figure 2b and Figure 2e).

**Figure 1.**
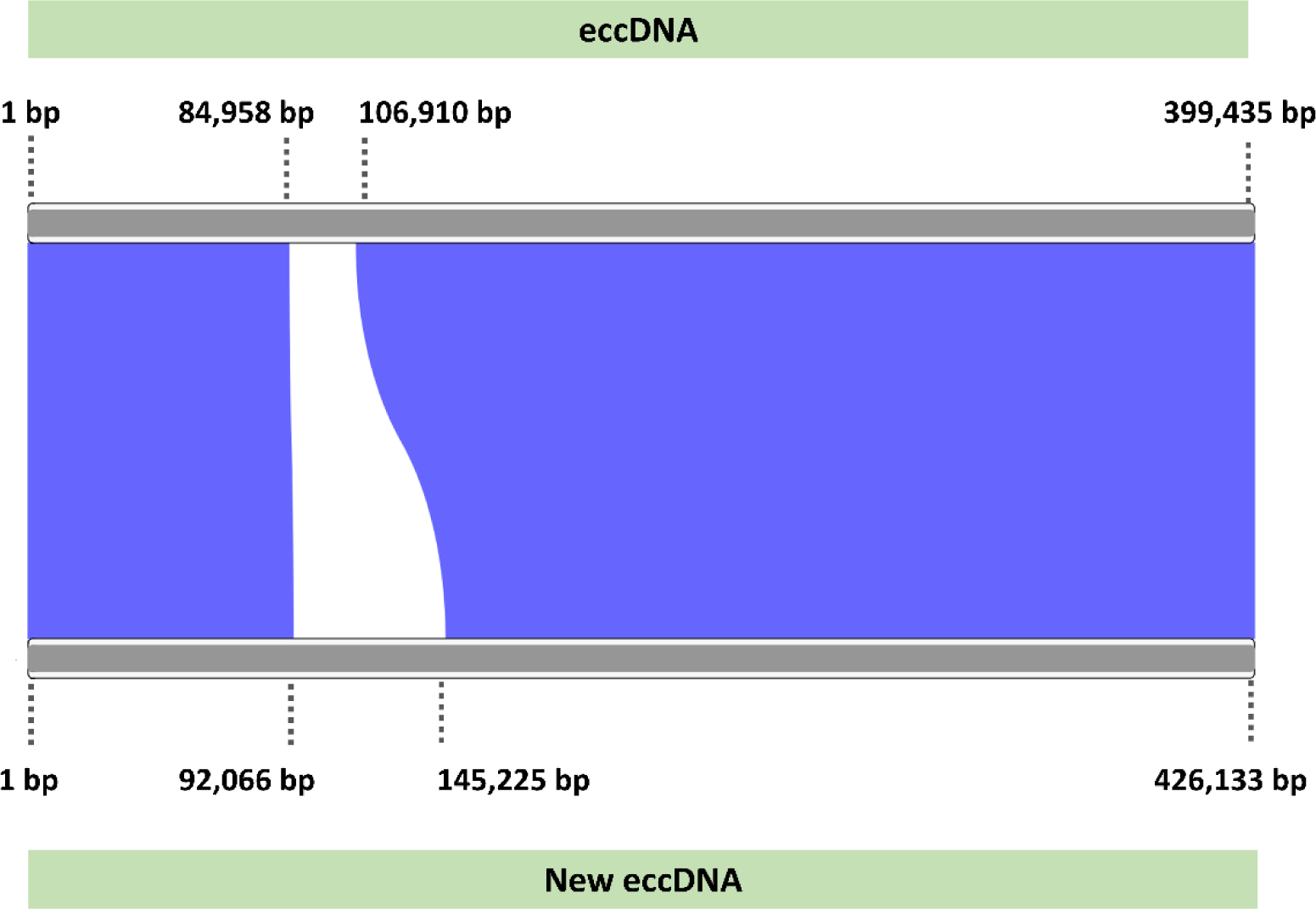
Comparison of the previously published A. palmeri eccDNA sequence from A. palmeri resistant to glyphosate (top) with the new sequence identified in a glufosinate-ammonium-resistant accession (bottom). The blue band indicates shared synteny between the two sequences. The white band represents areas without synteny between the two eccDNAs. Source sequencing for the *EPSPS* eccDNA was obtained from NCBI accession MT025716 (Molin et al. 2020a).

**Figure 2.**
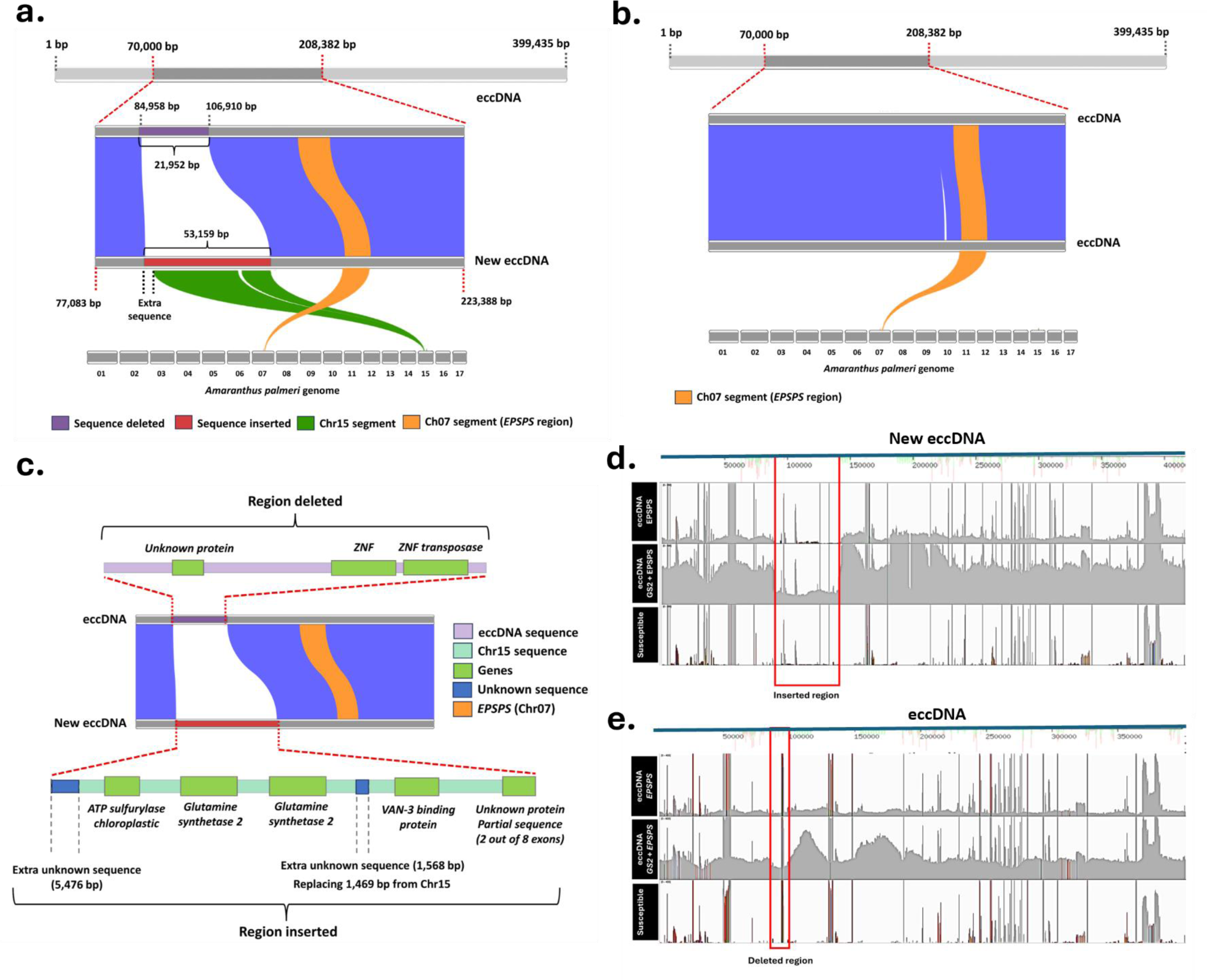
**(a)** Comparison of the previously published *Amaranthus palmeri* eccDNA sequence from *A. palmeri* resistant to glyphosate (top) with the new sequence identified in a glufosinate-ammonium-resistant accession (bottom) showing the size of the deleted and inserted fragments and the synteny of the insertion and *EPSPS* region with *A. palmeri* genome. **(b)** Comparison of the previously published A. palmeri eccDNA sequence from *A. palmeri* resistant to glyphosate (top) with the second eccDNA identified in the same glufosinate-ammonium-resistant accession (bottom) showing the EPSPS region synteny with A. palmeri genome. **(c)** Detailed composition of the sequence deleted from the original A. palmeri eccDNA (top) and the sequence inserted in the new eccDNA (bottom). **(d)** Reads from *A. palmeri* accession (MSR2) resistant to glyphosate, glyphosate/glufosinate-ammonium, and susceptible covering the new eccDNA, highlighting the new sequence inserted region. **(e)** Reads from *A. palmeri* accession resistant to glyphosate, glyphosate/glufosinate-ammonium, and susceptible covering the known eccDNA, highlighting the sequence absent in the new eccDNA.

The detailed visualization of the rearrangements shows coding genes deleted and inserted in the new eccDNA. The 21,952 bp fragment deleted had three genes encoding a gene of unknown function, a zinc finger (*ZFN*) containing gene, and a *ZFN transposase* gene (Figure 2c). The sequence replacing this locus has four complete protein-coding genes, ATP-sulfurylase chloroplastic (*APS*), two tandemly arranged glutamine synthetase 2 (*GS2*) genes, and a VAN-3 binding protein (*VAB*), as well as a truncated, partial gene of unknown function (two out of eight exons) (Figure 2c). This insert matches the *GS2* native location in chromosome 15 of the susceptible (Sus) *A. palmeri* genome (Chr15:9,240,010 bp – 9,283,115 bp). The presence of two *GS2* isoforms in tandem is also verified in its native location in chromosome 15. The contiguity of the chromosome 15 inserted fragment in the eccDNA is interrupted by a 1,568 bp of unknown sequence origin that replaces a 1,469 bp in the native location that was not inserted in the eccDNA.

By aligning raw PacBio reads to the two eccDNA assemblies, we were able to validate the assembly and support the *GS2* insertion. The new eccDNA only has continuous read alignments in the MSR2 accession (resistant to glufosinate-ammonium/glyphosate) (Figure 2d). One MSR2 individual resistant only to glyphosate (MSR2-Gly-R) shows read coverage only in regions of the new eccDNA that are identical to the original eccDNA and do not support the insertion. The original eccDNA shows long-read coverage across the whole assembled sequence in both individuals from MSR2 accession (Figure 2e). These results support the assemblies and the hypothesis that the glufosinate-ammonium + glyphosate-resistant carries two variants of the eccDNA, one carrying *GS2* + *EPSPS* genes and one with only *EPSPS*. The susceptible does not show any read coverage in both eccDNAs investigated except where they contain the native sequence carrying *GS2* and *EPSPS* as they are present in the susceptible genome.

When reads are aligned to the *A. palmeri* genome, the MSR2 (carrying the two types of eccDNA) shows a higher long-read coverage at both the *GS2* and *EPSPS* native location on chromosomes 15 and 7, respectively, when compared to the susceptible biotype (Figure 3a), indicating that these two regions are in a high copy eccDNA. The *ALS* gene was utilized as a reference gene, and no differences were observed in either accession. These results corroborate the higher copy number for the genes in that location and the eccDNA new insertion origin. The average read depth for the *ALS*, *EPSPS*, and *GS2* genes was calculated for the susceptible plants with no eccDNA present, plants with only *EPSPS* eccDNA (MSR2-Gly-R), and plants with both eccDNAs (glufosinate-ammonium/glyphosate-resistant). The MSR2 individual carrying both versions of eccDNA had six times the read depth for the *EPSPS* region (∼3×10^7^) when compared to MSR2-Gly-R with eccDNA carrying only *EPSPS* (∼5×10^6^), which indicates an additive copy number for *EPSPS* between the two replicons (Figure 3b). The increase in *EPSPS* copy number was expected due to the co-existence of two types of eccDNAs carrying this gene in a glufosinate-ammonium/glyphosate-resistant accession. Both accessions showed higher *EPSPS* read depth than the susceptible. Only the MSR2 carrying both versions of eccDNA showed higher read depth for the *GS2*, which was 11.5 times higher than both susceptible plants and those with only the original eccDNA (Figure 3b). No increase in read depth was observed for the *ALS* region in any of the accessions analyzed, indicating the absence of duplication of this region and suitability for using it as a reference.

**Figure 3.**
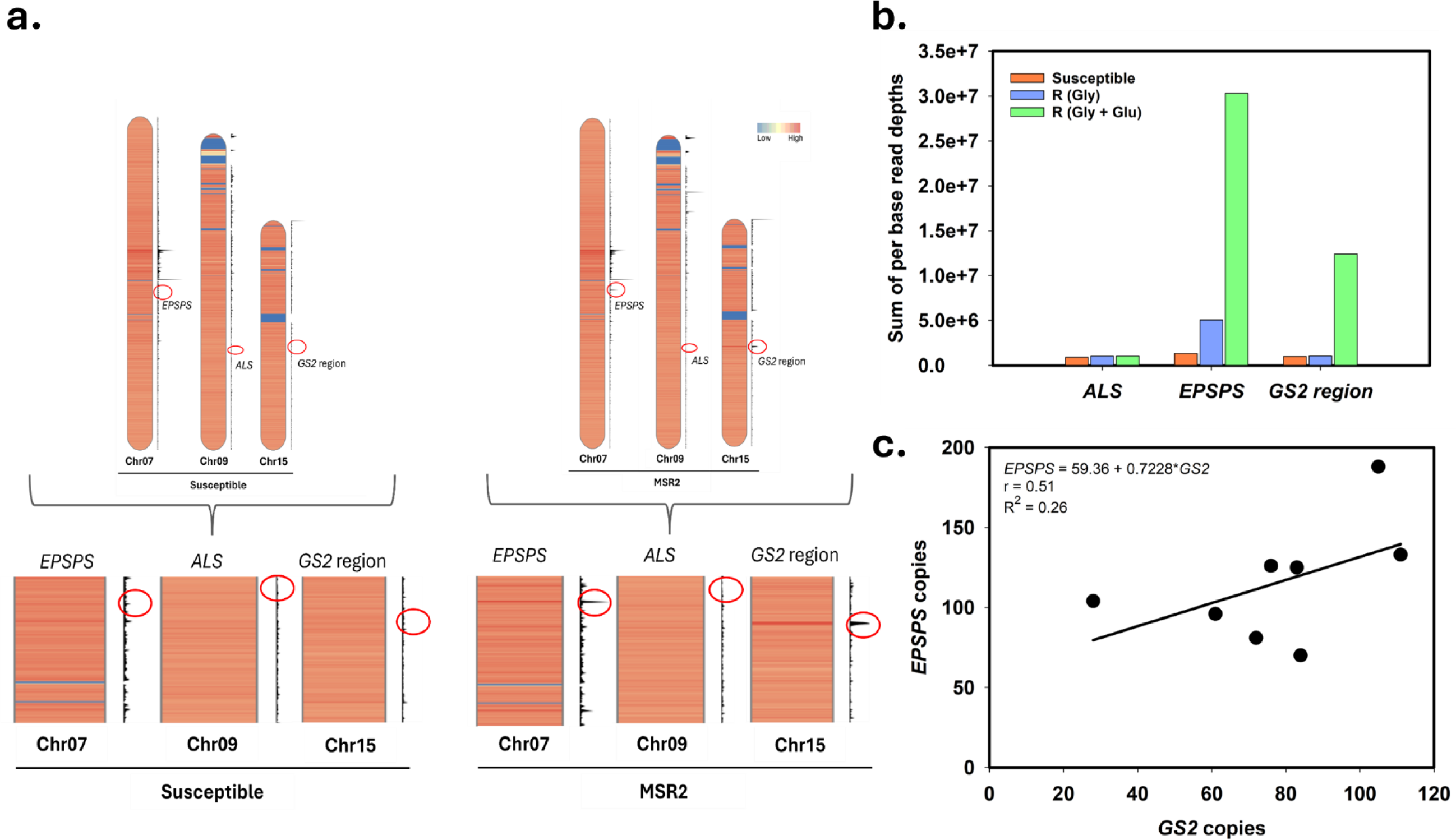
**(a)** Reads depth at *5-enolpyruvylshikimate-3-phosphate synthase* (*EPSPS*), *acetolactate synthase* (*ALS*), and *glutamine synthetase 2* (*GS2*) native locations between susceptible and glufosinate-ammonium/glyphosate-resistant plants. The *EPSPS*, *ALS*, and *GS2* regions are present in the *Amaranthus palmeri* chromosomes 07, 09, and 15, respectively. The color gradient from blue to red shows regions with low or high read depths, respectively. Red circles on the black scales next to each chromosome show the region of interest (*EPSPS*, *ALS*, or *GS2*). **(b)** Sum of per base read depths of *ALS*, *EPSPS*, and *GS2* genes in *Amaranthus palmeri* susceptible with no eccDNA (orange), MSR2 glyphosate-resistant with eccDNA carrying only *EPSPS* (blue), and MSR2 glufosinate-ammonium/glyphosate-resistant with two eccDNAs, one carrying *EPSPS* + *GS2* and one with only *EPSPS* (green). **(c)** Correlation of the number of *GS2* and *EPSPS* genes in eight *A. palmeri* plants from MSR2 accession resistant to glyphosate/glufosinate. *r,* correlation coefficient of Pearson’s correlation test.

The MSR2 carrying both eccDNA versions showed 2.5 times more read depth for *EPSPS* than *GS2*, further supporting the discovery that both replicons contain the *EPSPS* while only one contains the *GS2* genes; furthermore, since it is >2, this indicates the original replicon is present in a 1.5:1 ration original:new eccDNA ratio in the plants analyzed (Figure 3b). This eccDNA abundance difference observed between the *GS2* + *EPSPS* eccDNA and *EPSPS* eccDNA was also validated through the correlation of *EPSPS* × *GS2* copy number in additional *A. palmeri* plants from this accession. Seven out of eight plants showed higher *EPSPS* copies than *GS2* at various proportions, ranging from ∼1.13 to 3.71 times more (Figure 3c).

### Validation of the glufosinate-resistant eccDNA replicon assembly

Four primer sets were designed and tested to validate the glufosinate-resistant eccDNA replicon assembly (Figure 4A), and the PCR assays were performed comparing two susceptible, two glyphosate-(Gly-R), and four glufosinate-resistant (MSR2) individuals. Importantly, the MSR2 individuals used in this study were also resistant to glyphosate. The markers 1 (577 bp) and 2 (948 bp) were glufosinate-resistant specific, targeting the junctions of the new rearrangement in eccDNA whereas markers 3 (635 bp) and 4 (743 bp) were glyphosate-resistant specific, targeting the junctions of *EPSPS* insertion in the eccDNA and the original region that was deleted in the new eccDNA, respectively (Figure 4A and B). The PCR-based assays showed amplification of markers 1 and 2 in MSR2, which corroborates the assembly of the glufosinate-resistant eccDNA replicon as well as its occurrence in glufosinate-resistant individuals. The markers 3 and 4 (glyphosate-resistant specific) were present in both Gly-R and MSR2 populations (Figure 4B-3 and 4B-4), however, MSR2-plant 1 individual did not show this amplicon (Figure 4B-4). Susceptible individuals did not show amplification of either glufosinate-resistant- or glyphosate-resistant-specific markers.

**Figure 4.**
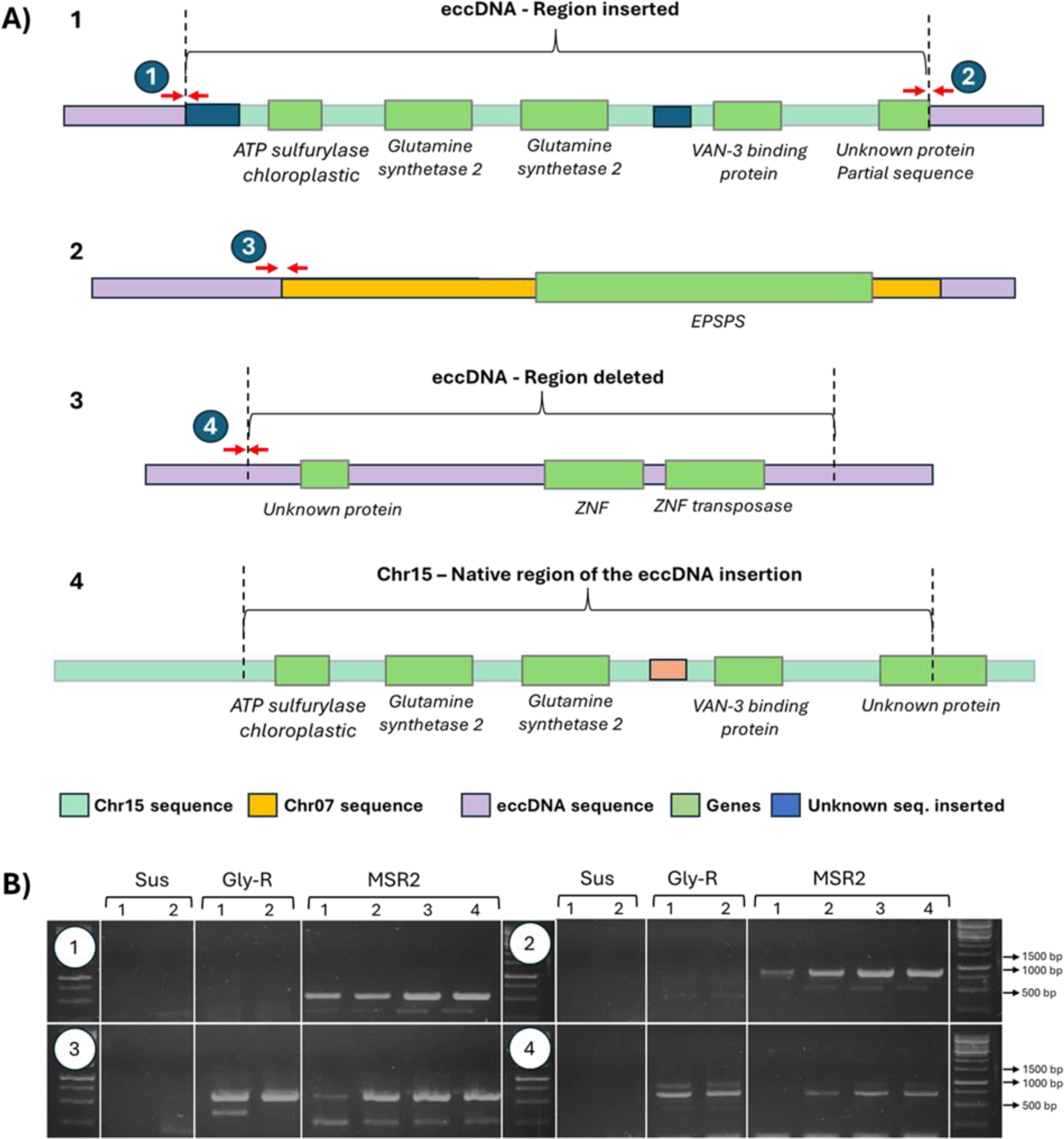
**(A)** The native locus region surrounding *GS2* (C), eccDNA-*EPSPS* (B), and eccDNA-*EPSPS* + *GS2* (A) in *Amaranthus palmeri*. Numerated junctions were used to validate model assembly through PCR-based molecular markers. **(B)** PCR that confirms the junctions are in MSR2 and not Sus. Also confirms that the MSR2 has EPSPS-only replicon and the EPSPS/GS2.1-GS2.2 replicon in the same plants.

### qPCR assay displayed copy number variations of different targeted genes associated with the new eccDNA rearrangement

The copy number of the genes occurring in the original and new eccDNA replicons were assessed in individuals from the susceptible (SUS), only glyphosate-resistant (Gly-R), and glufosinate + glyphosate-resistant (MSR2) accessions. All susceptible individuals displayed one *EPSPS* and two *GS2* copies, confirming the absence of the either eccDNA replicons (Supplementary Table 4). Similarly, these individuals also displayed one copy of *APS* and *VAB* genes. However, the unknown gene, *ZNF* and *ZNF* transposase genes copies were greater than one in most of susceptible individuals, indicating multiple copies in the genome. The Gly-R individuals displayed *EPSPS* copy number from 52 to 213 copies, and also higher copy number for *UKN*, *ZNF*, and *ZNF transposase* confirming the occurrence of the original eccDNA replicon carrying only the herbicide target *EPSPS*. However, the number of these genes did not perfectly correlate with the *EPSPS* copies. For example, Gly-R-1 individual showed 213 *EPSPS* while showing nearly half copies of the *UKN* gene present in the *EPSPS* replicon, and nearly one copy of the *ZNF* and *ZNF transposase* genes. In the meantime, Gly-R-2 individual showed 150 *EPSPS* copies and 99 copies of the *UKN* gene, while also showing 236 copies of the *ZNF* and 208 copies of the *ZNF* transposase genes. Importantly, all Gly-R individuals showed two *GS2* and one *APS* and *VAB* gene copies, indicating the absence of the new *EPSPS* + *GS2* eccDNA. The MSR2 individuals showed a range of 7 to 247 *GS2* copies, and also increased copy number for *APS* and *VAB* confirming the occurrence of the new *EPSPS* + *GS2* eccDNA in this accession. The number of *APS* and *VAB* genes were nearly half of the *GS2* for the majority of the individuals tested. However, MSR2-2 had 44 copies of *APS* while displaying 12 *VAB* gene and 24 *GS2*, suggesting the occurrence of structural variants among *EPSPS* + *GS2* eccDNA replicon. All MSR2 individuals also displayed increased *EPSPS* (47 to 182 copies), *UKN*, *ZNF*, and *ZNF transposase* copy number, confirming also the presence of the original eccDNA in the same plant, as assembled in our study. In time, MSR2 individuals also indicated the occurrence of variability in the *EPSPS* replicon. For example, MSR2-2 had 173 *EPSPS* copies, 105 copies of the *UKN* gene, while also showing 4 and 7 copies of the *ZNF* and *ZNF* transposase genes, respectively. On the other hand, MSR2-9 showed 182 *EPSPS* and 118 *UKN* gene copies, but 272 *ZNF* and 247 *ZNF* transposase copies (Supplementary Table 4).

### Origins of the eccDNA in *A. palmeri* genome

A synteny analysis comparing the new eccDNA with the entire *A. palmeri* genome was performed to identify the origins of the whole sequence in the new eccDNA. When analyzing small synteny blocks (> 2000 bp), it is possible to identify many eccDNA fragments that are syntenic to several loci in the genome, most of which contain predicted transposable elements (Figure 5). Surprisingly, a considerable portion of the eccDNA had similarity >1000bp with any sequence in the *A. palmeri* genome (Supplemental Figure S2). The results show that the new eccDNA was not formed in a straightforward way through wholesale excision from the genome. When the synteny analysis was restricted to blocks larger than 2000 bp, three genes were annotated, one encoding an F-box/LRR-repeat protein and two Aminotransferase-like plant mobile domain, syntenic to chromosomes 14 and 15, respectively, as well as the *APS*, *GS2* isoforms, *VAB* binding protein, and *EPSPS* described before (Figure 5). In addition, transposable elements with synteny to several chromosomes were annotated. Three long terminal repeats (LTRs) retrotransposons, also known as class I transposable elements, one *Copia*, one *Gypsy*, and one unknown family (Figure 5).

**Figure 5.**
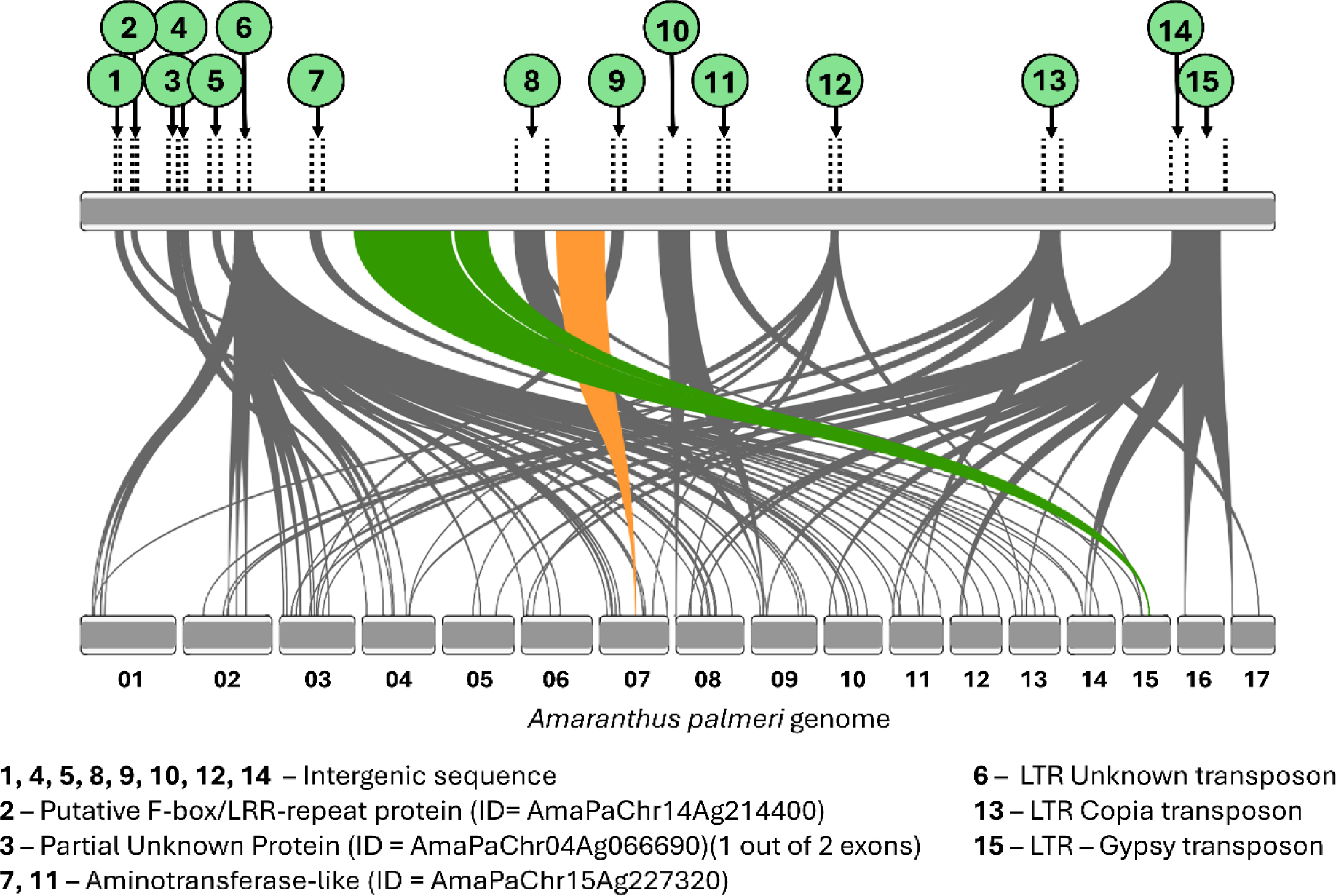
Annotation of synteny blocks bigger than 2000 bp between the new *EPSPS* + *GS2* eccDNA assembly (top row) and *Amaranthus palmeri* genome (bottom row). The green and orange lines show the syntenic blocks containing the *enolpyruvylshikimate-3-phosphate synthase* (*EPSPS*) and the chloroplastic *glutamine synthetase* (*GS2*) gene regions. Transposable elements were annotated using the EDTA pipeline.

Next, there was an effort to determine whether transposon activity could be implicated in the *GS2* insertion event. Transposable elements in the new eccDNA assembly originated from Chr07 and 15, and associated flanking regions were investigated. The segment deleted from the original eccDNA did not have any transposable element within or flanking the inserted sequence (Supplemental Figure S3). In the insertion containing *GS2* genes in the new eccDNA, there were three transposons, two hAT (MITEs) and one CACTA, both transposons class II (cut-and-paste) type TEs, located at the beginning of each side of the insertion (Supplemental Figure S3). No transposons were annotated in the 20,000 bp region flanking the new insertion. There were two transposable elements annotated in the insertion with *EPSPS* in both original and new eccDNA sequences, a Mutator and a Tc1/mariner, also class II transposons (Supplemental Figure S4). A hAT (MITE) was identified in the eccDNA sequence flanking the *EPSPS* insertion (Supplemental Figure S4). When analyzing the *GS2* insertion chromosomic native region in Chr15, only one hAT (MITE) transposon was identified downstream of the segment transferred to the eccDNA (Supplemental Figure S3). The *EPSPS* native region in Chr07 has a Tc1/mariner in the region inserted to the eccDNA, and a Mutator flanking the region (Supplemental Figure S3). When comparing the Tc1/Mariner and Mutator transposons associated with the *EPSPS* insertion in eccDNA and the Chr07 native region, they are located in different positions. In the native location in Chr07, the Mutator is outside of the sequence rearranged, while it is in the inserted sequence in the new eccDNA; the Tc1/mariner is up-stream of the *EPSPS* in the Chr07 native region, while it is downstream of *EPSPS* in the sequence inserted in the eccDNA (Supplemental Figure S5).

### Investigation of *glutamine synthetase* amplification in a second *A. palmeri* accession (MSR1)

A second *A. palmeri* accession called MSR1, also resistant to glufosinate-ammonium and glyphosate, was utilized to further investigate the *glutamine synthetase* gene amplification. Comparison of the copy number of *GS2* isoforms between MSR1 and MSR2 accessions showed completely different patterns. The MSR1 accession showed amplification of only the *GS2.1* isoform gene (0.90-34.07 copies) in addition to *EPSPS*, while MSR2 had amplification of both *GS2.1* (0.75-15.10 copies) and *GS2.2* (0.95-36.59 copies) isoforms (Figure 6). However, in MSR2 accession, the isoform *GS2.2* is more abundant than the *GS2.1*, 0.93-36 copies and 0.75-15 copies, respectively (Figure 6). This result suggests the presence of a third eccDNA containing only *GS2.2* in MSR2, which was not assembled in the present study, and potentially another eccDNA containing only *GS2.1* in MSR1. Amplification of the *EPSPS* gene was observed in both MSR1 and MSR2, ranging from 2- to 32-fold. The *GS1* isoforms were not amplified in either accession.

**Figure 6.**
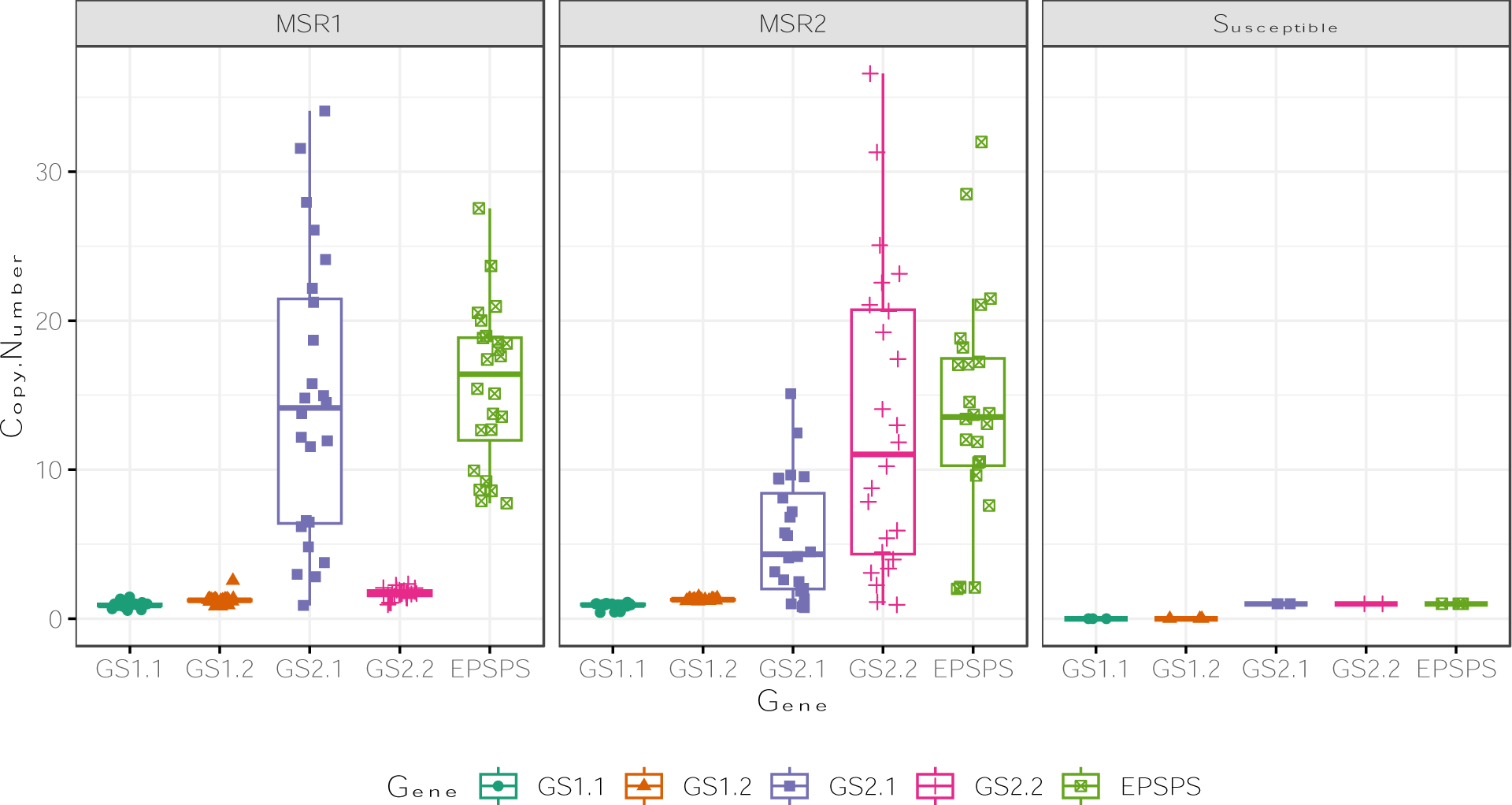
Box plots of the copy number of the four glutamine synthetase genes and one *EPSPS* gene in two glyphosate/glufosinate resistant *Amaranthus palmeri* accession MSR1, MSR2, and on glyphosate/glufosinate susceptible accession as estimated using digital PCR and RT-qPCR. Abbreviations: *GS1.1* and *GS1.2*, cytosolic glutamine synthetase isoforms; *GS2.1* and *GS2.2*, chloroplastic glutamine synthetase isoforms; *EPSPS*, 5-enolpyruvylshikimate-3-phosphate synthase. Box plots were generated using the packages “ggplot2” and “ggpubr” in RStudio (The R Project for Statistical Computing, Vienna, Austria).

Interestingly, when resistance gene copy number was estimated with digital PCR and then correlated, there is no consistent co-duplication of *GS2.1* and *EPSPS* in MSR1, implying the amplification of *GS2.1* in this accession is not on the same replicon or through the same mechanisms as *EPSPS* (Figure 7). It is also observed that an extremely high correlation between *GS2.1* and *GS2.2* (*r*=0.988) in MSR2, but at a ratio of ∼2:5 for *GS2.1:GS2.2,* indicating that *GS2.1* always duplicates with *GS2.2*; however, *GS2.2* often amplifies without *GS2.1*. This observation further supports a third, *GS2.2*-only variant of the eccDNA that has not yet been identified. Additionally, both *GS2.1* and *GS2.2* are significantly correlated with *EPSPS* amplification in MSR2, but it is a weaker correlation (*r*=0.72) compared to the *GS2* isoforms. This is most likely due to *EPSPS* independently changing the copy number of its original replicon without co-duplication of *GS2.1* or *GS2.2*.

**Figure 7.**
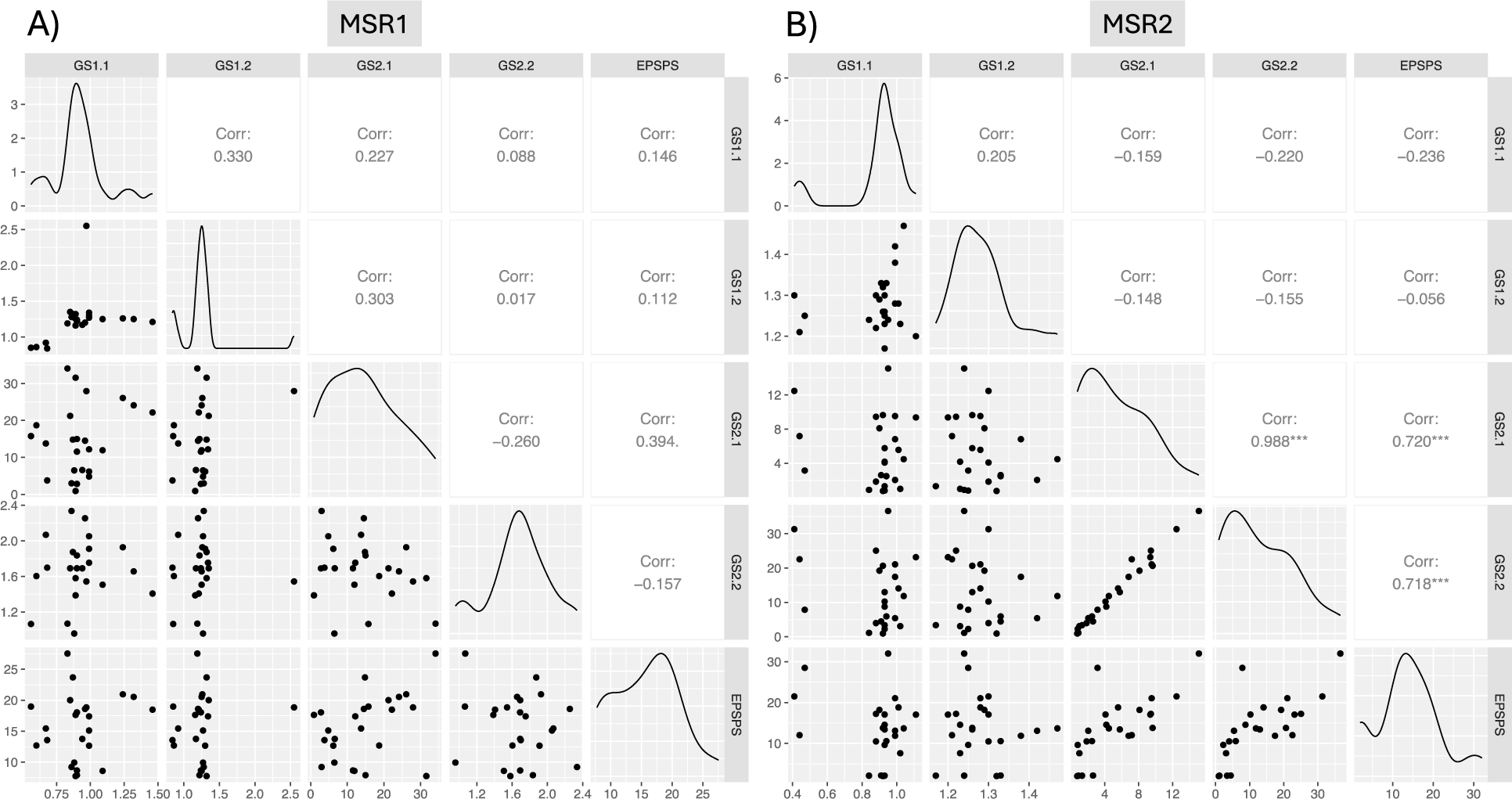
Linear correlation of the copy number between the four glutamine synthetase genes and one *EPSPS* gene in two glyphosate/glufosinate-resistant *Amaranthus palmeri* accessions MSR1 and MSR2. Correlation analysis was performed using the function “ggpair” under the package “GGally” in RStudio (The R Project for Statistical Computing, Vienna, Austria). Corr, correlation coefficient. ***, Significant at *p* < 0.001

## Discussion

Gene amplification and expression is an evolution mechanism associated with adaptative traits, such as resistance to antibiotics, pathogens, and pesticides (Almal and Padh 2012; Gaines et al. 2010; Picart-Picolo et al. 2020). Resistance to herbicides due to gene amplification was first reported in glyphosate-resistant *A. palmeri* (Gaines et al. 2010). Following this initial report, this mechanism has been observed in both monocotyledonous and eudicotyledonous species resistant to herbicides pertaining to different sites of action, such as glyphosate, glufosinate, and cyhalofop (Carvalho-Moore et al. 2022; Chen et al. 2015; Koo et al. 2023; Laforest et al. 2017). In glyphosate-resistant *A. palmeri*, eccDNAs carrying extra copies of the *EPSPS* gene have been detected and are directly associated with spreading herbicide resistance (Koo et al. 2018b; Molin et al. 2020a).

A glufosinate-resistant accession (MSR2) was previously characterized with a higher *GS2* copy number, without any *GS1* copy number changes, and no *GS1* and *GS2* mutations were detected (Carvalho-Moore et al. 2022). Since this accession also showed glyphosate resistance (Supplemental Figure S1), the initial hypothesis of this study was that gene amplification of *GS2* was also located in an eccDNA molecule, as reported for glyphosate resistance (Koo et al. 2018b). This hypothesis was confirmed by detecting a new eccDNA 426,133 long, including the two *GS2* isoforms and *EPSPS* genes co-occurring in the same eccDNA from the MSR2 accession. The assembly was validated by PCR amplification of the junctions and by detecting increased copy number of genes co-inserted with *GS2* isoforms. However, the same plant also carried the previously described *A. palmeri* eccDNA (*EPSPS* eccDNA) responsible for carrying only extra copies of the glyphosate target *EPSPS* (Koo et al. 2018b; Molin et al. 2020a). The new eccDNA is characterized by the excision of 21,952 bp and replaced by a 53,159 bp fragment from chromosome 15 compared to the first reported eccDNA. Except for this rearrangement, the two eccDNA versions are similar. Previous resequencing of divergent glyphosate-resistant *A. palmeri* populations showed that the shared eccDNAs are similar, without any structural variation (Molin et al. 2018). The fragment deleted from the eccDNA included regions encoding unknown protein (UNK), a zinc finger (*ZFN*), and a *ZFN transposase*, which can induce targeted double-strand DNA breaks and movement of transposable elements (Kim et al. 1996; Rice and Baker 2001). The mechanism of detaching and substituting the DNA fragment by the chromosome 15 region is unknown. Individuals carrying *EPSPS* + *GS2* eccDNA were likely randomly selected through frequent use of glufosinate. It is important to note that the native *GS2* sequence was still present in its innate location in chromosome 15 of the MSR2 containing *EPSPS* + *GS2* eccDNA, which would indicate DNA transposition, specifically copy-and-paste transposition mediated by retrotransposons, resulting in increased gene copy number (Hickman and Dyda 2016). The genes *APS, VAB,* and partial gene encoding an unknown protein were also co-duplicated with the two *GS2* isoforms in the new eccDNA. The effect of additional gene copies encoding *APS* and *VAB* still must be investigated. The new eccDNA containing *EPSPS* + *GS2* was completely assembled, and besides the *GS2* isoforms and *EPSPS*, no other genes encoding herbicide-targeted enzymes were inserted into the new replicon. Therefore, further investigations will be needed to determine why and how the *GS2* insertion occurred.

Even with the co-occurrence of two eccDNA versions, MSR2 showed different proportions of both versions; usually, the one with only *EPSPS* is more abundant. This result could indicate a high variation in how eccDNAs are transmitted across glufosinate-resistant individuals, which was, in some way, expected. According to Koo et al. (2018b), there are somatic variations among cells in glyphosate-resistant *A. palmeri* carrying the *EPSPS* eccDNA. In our studies, different MSR2 individuals showed a matching and mismatching copy number of targeted genes present in the *EPSPS* and *EPSPS* + *GS2* replicons, corroborating this finding. Koo et al. (2018) also found that the *EPSPS* eccDNA is not integrated into chromosomes and follows non-Mendelian segregation in gametes during meiosis, ultimately leading to cells with multiple or no *EPSPS* eccDNA. Individuals with higher gene copy number will have an undeniable survival advantage under herbicide selection pressure. This variability has been confirmed in previous works in which *EPSPS* copy number segregation in progenies carrying *EPSPS* eccDNA was random and unpredictable (Chandi et al. 2012; Giacomini et al. 2019; Mohseni-Moghadam et al. 2013).

Neither the new *EPSPS* + *GS2* nor the original *EPSPS* eccDNA were detected in the susceptible plants used as control. However, the existence of different and smaller eccDNAs sequences natively occurring have been detected in susceptible plants (Fu et al. 2023; Wang et al. 2021; Zhuang et al. 2024). In *A. palmeri*, eccDNAs ranging from 27 bp to 27,000 bp were identified in glyphosate-susceptible and resistant populations (Spier Camposano et al. 2022). Additionally, the eccDNA abundance might change under biotic or abiotic stress (Turner et al. 2017; Wang et al. 2021; Zhuang et al. 2024). For instance, a significantly higher amount of eccDNAs were quantified in rice leaves under darkness or UV-irradiation treatments compared to control plants maintained under normal light conditions (Zhuang et al. 2024). Most of the eccDNAs have the characteristic of rapid degradation. However, the ones that have the ability to amplify themselves and provide an adaptive advantage can survive (Zuo et al. 2022).

The presence of different *GS* isoforms was previously detected in *A. palmeri* accession from Missouri and *L. perenne* spp. *multiflorum* from Oregon (Brunharo et al. 2019; Noguera et al. 2022). In both species, the *GS1* isoforms did not show differences in amplification or expression. Like what was observed in the glufosinate-resistant accession MSR1, the *A. palmeri* accession from Missouri, discussed by Noguera et al. (2019), showed increased amplification of only the *GS2.1* isoform with no *GS2.2* amplification. In contrast, MSR2 individuals showed amplification of both isoforms, which supports the existence of a replicon with the arrangement carrying *EPSPS* + *GS2*. Moreover, MSR2 showed a much higher abundance of *GS2.2* compared to *GS2.1*. These results indicate the existence of eccDNAs carrying only one isoform type or carrying both and a different co-evolution pattern for different locations. Circular DNAs of different sizes were identified in herbicide-susceptible and resistant plant populations, which could serve as a font of genetic diversity during evolution with the possibility of different replicon structures (Spier Camposano et al. 2022; Fu et al. 2023).

The identification of *EPSPS* amplification as a resistance mechanism, followed by the discovery of eccDNAs carrying it (Gaines et al. 2012; Koo et al. 2018b; Molin et al. 2020b), changed weed scientists’ perception of herbicide resistance evolution. Due to the eccDNA’s autonomous replication, cross-pollination characteristic of *A. palmeri,* and small seeds favorable to dissemination, glyphosate resistance due to *EPSPS* amplification is widely spread in *A. palmeri* accessions across the world. The findings of this study showed that *GS2* was also inserted in eccDNAs, and a quick spread of resistance is expected due to the *A. palmeri* characteristics, as verified for glyphosate. The eccDNA possesses the ability to aggregate and amplify multiple genes encoding herbicide targets, as shown in this study, but also genes associated with herbicide resistance mechanisms, like *GSTF1* in *A. myosuroides* (Fu et al. 2023). Consequently, the progeny carrying the eccDNAs accumulating resistance to distinct herbicide sites of action will undoubtedly have an advantage under different selection pressure scenarios. Our findings show the ability of *A. palmeri* to collect and accumulate extra pieces of DNA in existent replicons, displaying the complexity of eccDNAs and their importance to weed evolution dynamics. Rapid adaptation to different herbicides is also a threat to individuals carrying eccDNAs due to the inherent possibility of this event occurring again, incorporating other herbicide targets. Glyphosate and glufosinate are major allies in the current practices used in food production, and a single event driving the resistance to both herbicides has massive impacts on agriculture.

## Experimental Procedures

### Plant material

A glufosinate-resistant *A. palmeri* accession from Arkansas (MSR2) confirmed to show high resistance levels (R/S ratio: 24-fold; Priess et al. 2022) was selected for this study. The resistant accession was sprayed with a 4× glufosinate dose (1× labeled rate = 655 g ai ha^−1^), and survivors were cross-pollinated. Importantly, this accession was also confirmed resistant to glyphosate (Supplementary Figure S1). Additionally, one susceptible accession collected in 1986 was used for comparison. The MSR2 and susceptible seeds were sown into 7-cm diameter plastic pots filled with potting mix (Sun Gro^®^ Horticulture, Agawam, MA, USA). Seedlings were grown in a greenhouse at 25 – 30 °C, with a 16 h/8 h day/night period at the Milo J. Shult Agricultural Research and Extension Center in Fayetteville, AR, USA. Leaf tissue from three MSR2 (*n=* 3) and three susceptible (*n=* 3) untreated seedlings were used for genomic high molecular weight DNA isolation using the Quick DNA HMW MagBead Kit (Zymo Research, Irvine, CA, USA). DNA integrity, concentration, and quality were verified, and the average fragment size was above 40 kb.

### Sequencing and assembly of Amaranthus palmeri accessions

High molecular weight DNA of three MSR2 and three susceptible individuals were submitted to whole genome sequencing by Pacific Biosciences (PacBio) Revio™ system with high fidelity (HiFi) library preparation. The output average of bases per sample and read length were 12.16 gbp and 15.29 kbp for MSR2, and 13.48 gbp and 16.61 kbp for susceptible. The eccDNA was assembled utilizing *flye v. 2.9.2* assembler (Kolmogorov et al. 2019) and curated manually utilizing the PacBio HiFi long reads and the published version of the eccDNA (NCBI accession: MT025716; Molin et al. 2020a). The manual curation involved the validation of the eccDNA contigs built by *flye* and connecting different contigs, utilizing BLAST+ v.2.5.0 (Altschul et al. 1990), SAMtools v1.18 (Li et al. 2009), Minimap2 v2.17 (Li 2018), RagTag v2.1.0 (Alonge et al. 2022), and Mummer3 v. 3.23 (Kurtz et al. 2004) to identify and align the regions of similarity between them. After completion, the complete assemblies with long-reads supporting were visualized in the Integrative Genomics Viewer (IGV) v2.11.0 (Robinson et al. 2011). Due to segregation, we noticed one out of three MSR2 individuals sequenced only one had the new eccDNA carrying *GS2* isoforms + *EPSPS* (utilized to assemble the new eccDNA in our study), one had only the original eccDNA carrying only *EPSPS*, and a third did not have either eccDNA, probably susceptible. We cannot discard the hypothesis that the MSR2 individual characterized with only the original *EPSPS* eccDNA has a different eccDNA not assembled in our study carrying one or more *GS2* isoforms.

### Synteny analyzes

Syntenic analyses were generated utilizing Minimap2 v2.17 (Li 2018), RagTag v2.1.0 (Alonge et al. 2022), and Mummer3 v. 3.23 (Kurtz et al. 2004). RIdeogram package (Hao et al. 2020) in R was utilized to generate the images.

### PCR-based validation of glufosinate-resistant eccDNA replicon in Amaranthus palmeri

PCR-based markers were developed to validate the assembly of the glufosinate-resistant eccDNA replicon. Primers were designed to amplify fragments that include the junctions of the region inserted in the glufosinate-resistant eccDNA replicon (markers 1 and 2, Figure 4A) and deleted from the glyphosate-resistant eccDNA replicon (markers 3 and 4, Figure 4A). DNA from two susceptible and four MSR2 individuals were used in this assay. Also, DNA from two glyphosate-resistant (Gly-R) individuals, previously characterized (Sulzback et al. 2024) were included. Approximately 100 mg of young leaf tissue was ground using a plastic pestle in a conic 1.5 mL microcentrifuge tube to obtain a fine powder. High molecular weight DNA was extracted utilizing Wizard^®^ HMW DNA Extraction Kit (Promega Corporation, Madison, WI, USA) following the manufacturer’s protocol. The DNA obtained was diluted to a final concentration of 50 ng µL^−1^ for further use. The reactions were performed with 2 µL of DNA (50 ng µL^−1^), 1 µL of forward and 1 µL of reverse primers (10 mM), 1 µL of MgCl_2_ (25 mM), 7.5 µL of nuclease-free water, and 12.5 µL of GoTaq^®^ G2 Green Master Mix (Promega), with a total of 25 µL per reaction. The thermal cycler settings were as follows: 95 °C for 5 mins, and 95°C for 30 secs, annealing temperature for 30 secs, 72 °C during the extension time repeated for 40 cycles, and a final extension at 72 °C for 10 mins. Primers sequences, annealing temperature, extension time, and the length of the amplicons are described in Supplementary Table 1. The presence of amplification was observed in agarose gel (1.5%) (Figure 4B).

### qPCR-based validation of glufosinate-resistant eccDNA replicon in Amaranthus palmeri

A copy number variation assay was also developed to support the new glufosinate-resistant eccDNA replicon assembly. In this assay, the number of ATP-sulfurylase chloroplastic (*APS*), VAN-3 binding protein (*VAB*), and of that unknown (*UKN*) partial sequence genes (Figure 4A) should correlate to glutamine synthetase 2 (*GS2)* in glufosinate-resistant individuals, since these genes are co-occurring in the new eccDNA replicon assembled; primer targeting *GS2* were binding both isoforms *GS2.1* and *GS2.2* Similarly, the number of *EPSPS*, *UNK*, *ZNF* and *ZNF*-transposase genes should correlate in DNA from glyphosate- and glufosinate + glyphosate-resistant individuals. Primers (Supplementary Table 2) were designed to amplify the different targets in the eccDNA assembled in this study. The reactions were performed with 2 µL of DNA (15 ng µL^−1^), 1 µL of forward and 1 µL of reverse primers (10 mM), 6 µL of nuclease-free water, and 10 µL of SsoAdvanced^™^ Universal SYBR^®^ Green Supermix (Bio-Rad), with a total of 20 µL per reaction. The real-time qPCR cycles started with 3 min at 95 °C and were followed by 39 cycles at 95 °C for 10 secs and 60 °C for 30 secs, with standard melting curves included in a CFX 96 Real-time system thermocycler (Bio-Rad). The assay was performed with three technical replicates per sample tested. The copy number of targeted genes was estimated relative to *ALS* in each biological replicate utilizing the method 2^−ΔCt^. The average gene copy number of each targeted gene was compared across biological replicates by Tukey HSD (α=0.05).

### Copy number of GS isoforms and EPSPS in different glufosinate-resistant Amaranthus palmeri accession

Besides MSR2, a different glufosinate-resistant accession (MSR1), also showing glufosinate-ammonium/glyphosate resistance and *GS2* amplification was identified in Arkansas (Priess et al. 2022). Assays were conducted to verify if the new replicon arrangement found in MSR2 was present in MSR1 plants. Additionally, the gene copy number variation of each *GS* isoform and *EPSPS* was quantified for MSR1 and MSR2. Susceptible plants were also included for comparison. For the preparation of the DNA, twenty-four leaf samples (0.5 cm²) of individual plants from each *A. palmeri* accession were transferred into a sample tube (collection microtubes; Qiagen, Hilden, Germany). The samples were then homogenized in a shaker mill (TissueLyser II; Qiagen, Hilden, Germany) with steel beads. DNA extraction was performed in the KingFisher^TM^ Flex Magnetic Particle Processors (Thermo Fisher Scientific, Schwerte, Germany) using the Chemagic Plant 400 kit (Perkin Elmer, Rodgau, Germany) according to the manufacturer’s instructions.

Nanowell-based digital PCR (dPCR) was performed to determine the CNV of *GS1.1* and *GS1.2* isoforms and *EPSPS*. The dPCR was conducted in a final volume of 12 µl using 1.48 µl of DNA, 0.48 µl (0.2 µM) of specific primers (Supplementary Table 3) and 0.25 µl (0.2 µM) of probe (biomers.net GmbH, Ulm, Germany), 3 µl of QIAcuity Probe PCR Kit mix, and 3.92 µl PCR-Grade H_2_O for the triplex dPCR. The assay was performed in a dPCR thermal cycler (QIAcuity One, 5plex Device, Qiagen, Hilden Germany) in a 96-well nanoplate with 8.500 nanowells for each sample (QIAcuity Nanoplate 8.5k 96-well) under the following conditions: 2 min at 95 °C and 55 cycles of 15 s denaturation at 95 °C; 40 s annealing, elongation, and detection at 60 °C. Partitions were imaged with the following conditions. FAM and HEX, 500 ms exposure time, gain set to 6; ROX 400 ms exposure time, gain set to 6. Qiagen’s QIAcuity Software Suite (version 2.1.8) was used to determine sample thresholds using positive, negative, and no-template control wells, and the copy number variation.

TaqMan^™^ technology was used to determine the gene copy number and expression of the *GS2* and *GS2.1* isoforms. The TaqMan^™^ assays were designed to allow a multiplex approach for the target and reference genes (Supplementary Table 3). The real-time quantitative PCR (RT-qPCR) was performed in a final volume of 25 µl using 5 µl of DNA, 1 µl (0.2 µM) of specific primers and 0.25 µl (0.2 µM) of probe (biomers.net GmbH, Ulm, Germany), 0.25µl of SNP PolTaq DNA Polymerase and 2.5 µl 10× amplification buffer (Genaxxon bioscience GmbH, Ulm, Germany), 0.5 µL (10mM) dNTP mix and 11.75 µl PCR-Grade H_2_O for the triplex RT-qPCR (*GS2.1*, *GS2.2* and *Actin*). Three technical replicates were used for each sample. RT-qPCR was performed in a qPCR thermal cycler (CFX96 Touch Real-Time PCR Detection System, Bio-Rad Laboratories GmbH, Germany) under the following conditions: 5 min at 95 °C and 35 cycles of 10 s denaturation at 95 °C; 30 s annealing, elongation, and detection at 60 °C. The evaluation, according to the 2^−ΔΔ*CT*^ method, was carried out with the software Bio-Rad CFX Maestro 2.2 version 5.2.008.0222.

### Genes and transposable elements annotation

The eccDNA sequences bigger than 2000 bp with synteny to the *A. palmeri* genome had the coding genes annotated based on the annotation file of the *A. palmeri* genome from the International Weed Genomics Consortium (IWGC) (Montgomery et al. 2024; Raiyemo et al. 2024). A synteny was performed between the eccDNA and the genome, and the coordinates were compared to the *A. palmeri* annotation file, checking the genes in the coordinates interval. The transposable elements were annotated utilizing Extensive de-novo TE Annotator (EDTA) pipeline (Ou et al. 2019).

### Statistical Analysis

Data visualization and statistical analyses were performed with JMP Pro 18 (SAS Institute Inc., Cary, NC), including Student’s t-test and ANOVA followed by Tukey HSD test, or RStudio using the packages “GGally”, “ggplot2”, and “ggpubr”. Significant differences were determined at *p* < 0.05.

### Accession numbers

All the data has been deposited in the NCBI database archive under the Bioproject accession number PRJNA1151833. The PacBio HiFi reads were deposited in the Sequence Read Archive (SRA) for susceptible individuals (SRR30359583, SRR30359584, and SRR30359585), the MSR2 individual resistant to glufosinate + glyphosate (SRR30359587), the MSR2 individual resistant to glyphosate (SRR30359588), and the MSR2 individual susceptible to glyphosate and glufosinate (SRR30359586). The new eccDNA assembly has been deposited under the accession number PQ252370.

## Acknowledgments

The authors thank the University of Arkansas Division of Agriculture and the Weed Science group for their support in this research.

## Author contributions

Conceptualization: PC-M, JKN, ELP, AP, IM, and JL; Project administration: PC-M, JKN, and ELP; EccDNA assembly: LC, PC-M, and ELP; Genomic analysis: LC; Molecular analyses and primer design: EAB, LC, and AP; Funding: AP, IM, and JL; Original draft: PC-M, LC, EAB, ELP, and JKN; Final review: all.

## References

Almal SH, Padh H. Implications of gene copy-number variation in health and diseases. J Hum Genet. 2012:57(1):6–13. 10.1038/jhg.2011.108

Alonge M, Lebeigle L, Kirsche M, Jenike K, Ou S, Aganezov S, Wang X, Lippman ZB, Schatz MC. Automated assembly scaffolding using RagTag elevates a new tomato system for high-throughput genome editing. Genome Biol. 2022:23(1):258. 10.1186/s13059-022-02823-7

Altschul SF, Gish W, Miller W, Myers EW, Lipman DJ. Basic local alignment search tool. J Mol Biol. 1990:215(3):403–410. 10.1016/S0022-2836(05)80360-2

Bayer E, Gugel KH, Hägele K, Hagenmaier H, Jessipow S, König WA, Zähner H. Stoffwechselprodukte von Mikroorganismen. 98. Mitteilung. Phosphinothricin und Phosphinothricyl-Alanyl-Alanin. Helv Chim Acta. 1972:55(1):224–239. 10.1002/hlca.19720550126

Brunharo CA, Takano HK, Mallory-Smith CA, Dayan FE, Hanson BD. Role of glutamine synthetase isogenes and herbicide metabolism in the mechanism of resistance to glufosinate in *Lolium perenne* L. spp. multiflorum biotypes from Oregon. J Agric Food Chem. 2019:67(31):8431–8440. 10.1021/acs.jafc.9b01392

Carvalho-Moore P, Norsworthy JK, González-Torralva F, Hwang J-I, Patel JD, Barber LT, Butts TR, McElroy JS. Unraveling the mechanism of resistance in a glufosinate-resistant Palmer amaranth (*Amaranthus palmeri*) accession. Weed Sci. 2022:70(4):370–379. 10.1017/wsc.2022.31

Chandi A, Milla-Lewis SR, Giacomini D, Westra P, Preston C, Jordan DL, York AC, Burton JD, Whitaker JR. Inheritance of evolved glyphosate resistance in a North Carolina Palmer amaranth (*Amaranthus palmeri*) biotype. Int J Agron. 2012(1):176108. 10.1155/2012/176108

Chen J, Huang H, Zhang C, Wei S, Huang Z, Chen J, Wang X. Mutations and amplification of EPSPS gene confer resistance to glyphosate in goosegrass (*Eleusine indica*). Planta. 2015:242:859–868. 10.1007/s00425-015-2324-2

Dillon A, Varanasi VK, Danilova TV, Koo D-H, Nakka S, Peterson DE, Tranel PJ, Friebe B, Gill BS, Jugulam M. Physical mapping of amplified copies of the 5-enolpyruvylshikimate-3-phosphate synthase gene in glyphosate-resistant *Amaranthus tuberculatus*. Plant Physiol. 2017:173(2):1226–1234. 10.1104/pp.16.01427

Fu W, MacGregor DR, Comont D, Saski CA. Sequence characterization of extra-chromosomal circular DNA content in multiple blackgrass (*Alopecurus myosuroides*) populations. Genes. 2023:14(10):1905. 10.3390/genes14101905

Gaines TA, Zhang W, Wang D, Bukun B, Chisholm ST, Shaner DL, Nissen SJ, Patzoldt WL, Tranel PJ, Culpepper AS, et al. Gene amplification confers glyphosate resistance in *Amaranthus palmeri*. Proc Natl Acad Sci. 2010:107(3):1029–1034. 10.1073/pnas.0906649107

Giacomini DA, Westra P, Ward SM. (2019) Variable inheritance of amplified EPSPS gene copies in glyphosate-resistant Palmer amaranth (*Amaranthus palmeri*). Weed Sci. 2019:67(2):176–182. 10.1017/wsc.2018.65

Hall N, Chen J, Saski C, Westra P, Gaines T, Patterson EL. FHY3/FAR1 transposable elements generate adaptive genetic variation in the *Bassia scoparia* genome. bioRxiv. 2023:2023–05. 10.1101/2023.05.26.542497

Hao Z, Lv D, Ge Y, Shi J, Weijers D, Yu G, Chen J. RIdeogram: drawing SVG graphics to visualize and map genome-wide data on the idiograms. PeerJ Comput Sci. 2020:6:e251. 10.7717/peerj-cs.251

Hickman AB, Dyda F. (2016). DNA transposition at work. Chem Rev. 2016:116(20):12758–12784. 10.1021/acs.chemrev.6b00003

Hoerlein G. Glufosinate (phosphinothricin), a natural amino acid with unexpected herbicidal properties. Rev Environ Contam Toxicol. 1994:138:73–147. 10.1007/978-1-4612-2672-7

Hotta Y, Bassel A. Molecular size and circularity of DNA in cells of mammals and higher plants. Proc Natl Acad Sci. 1965:53(2):356–362. 10.1073/pnas.53.2.356

Hull RM, King M, Pizza G, Krueger F, Vergara X, Houseley J. Transcription-induced formation of extrachromosomal DNA during yeast ageing. PLoS Biol. 2019:17(12):e3000471. 10.1371/journal.pbio.3000471

Hwang JI, Norsworthy JK, Piveta LB, Souza MCCR, Barber LT, Butts TR. Metabolism of 2,4-D in resistant *Amaranthus palmeri* S. Wats. (Palmer amaranth). Crop Prot. 2023:165:106169. 10.1016/j.cropro.2022.106169

Jugulam M, Niehues K, Godar AS, Koo D-H, Danilova TV, Friebe B, Sehgal S, Varanasi VK, Wiersma A, Westra P, et al. Tandem amplification of a chromosomal segment harboring *5-enolpyruvylshikimate-3-phosphate synthase* locus confers glyphosate resistance in *Kochia scoparia*. Plant Physiol. 2014:166(3):1200–1207. 10.1104/pp.114.242826

Kim YG, Cha J, Chandrasegaran S. Hybrid restriction enzymes: zinc finger fusions to Fok I cleavage domain. Proc Natl Acad Sci. 1996:93(3):1156–1160. 10.1073/pnas.93.3.1156

Kolmogorov M, Yuan J, Lin Y, Pevzner PA. Assembly of long, error-prone reads using repeat graphs. Nat Biotechnol. 2019:37(5):540–546. 10.1038/s41587-019-0072-8

Koo D-H, Ju Y, Putta K, Sathishraj R, Roma-Burgos N, Jugulam M, Friebe B, Gill BS. Extrachromosomal DNA-mediated glyphosate resistance in Italian ryegrass. Pest Manag Sci. 2023:79(11):4290–4294. 10.1002/ps.7626

Koo D-H, Jugulam M, Putta K, Cuvaca IB, Peterson DE, Currie RS, Friebe B, Gill BS. Gene duplication and aneuploidy trigger rapid evolution of herbicide resistance in common waterhemp. Plant Physiol. 2018a:176(3):1932–1938. 10.1104/pp.17.01668

Koo D-H, Molin WT, Saski CA, Jiang J, Putta K, Jugulam M, Friebe B, Gill BS. Extrachromosomal circular DNA-based amplification and transmission of herbicide resistance in crop weed *Amaranthus palmeri*. Proc Natl Acad Sci. 2018b:115(13):3332–3337. 10.1073/pnas.1719354115

Kurtz S, Phillippy A, Delcher AL, Smoot M, Shumway M, Antonescu C, Salzberg SL. Versatile and open software for comparing large genomes. Genome Biol. 2004:5:1–9. 10.1186/gb-2004-5-2-r12

Küpper A, Borgato EA, Patterson EL, Gonçalves Netto A, Nicolai M, Carvalho SJP, Nissen SJ, Gaines TA, Christoffoleti PJ. Multiple resistance to glyphosate and acetolactate synthase inhibitors in Palmer amaranth (*Amaranthus palmeri*) identified in Brazil. Weed Sci. 2017:65(3):317–326. 10.1017/wsc.2017.1

Laforest M, Soufiane B, Simard MJ, Obeid K, Page E, Nurse RE. Acetyl-CoA carboxylase overexpression in herbicide-resistant large crabgrass (*Digitaria sanguinalis*). Pest Manag Sci. 2017:73(11):2227–2235. 10.1002/ps.4675

Li H. Minimap2: pairwise alignment for nucleotide sequences. Bioinf. 2018:34(18):3094–3100. 10.1093/bioinformatics/bty191

Li H, Handsaker B, Wysoker A, Fennell T, Ruan J, Homer N, Marth G, Abecasis G, Durbin R, 1000 Genome Project Data Processing Subgroup. The sequence alignment/map format and SAMtools. Bioinf. 2009:25(16):2078–2079. 10.1093/bioinformatics/btp352

Luo J, Li Y, Zhang T, Xv T, Chen C, Li M, Qui Q, Song Y, Wan S. Extrachromosomal circular DNA in cancer drug resistance and its potential clinical implications. Front Oncol. 2023:12:1092705. 10.3389/fonc.2022.1092705

Massinga RA, Currie RS, Horak MJ, Boyer JE. Interference of Palmer amaranth in corn. Weed Sci. 2001:49(2):202–208. 10.1614/0043-1745(2001)049[0202:IOPAIC]2.0.CO;2

McNally SF, Hirel B, Stewart GR. Nitrogen-metabolism in halophytes. v. The occurrence of multiple forms of glutamine-synthetase in leaf tissue. New Phytol. 1983:94(1):47–56. 10.1111/j.1469-8137.1983.tb02720.x

Mohseni-Moghadam M, Schroeder J, Ashigh J. Mechanism of resistance and inheritance in glyphosate resistant Palmer amaranth (*Amaranthus palmeri*) populations from New Mexico, USA. Weed Sci. 2013:61(4):517–525. 10.1614/WS-D-13-00028.1

Molin WT, Yaguchi A, Blenner M, Saski CA. The eccDNA replicon: A heritable, extranuclear vehicle that enables gene amplification and glyphosate resistance in *Amaranthus palmeri*. The Plant Cell. 2020a:32(7):2132–2140. 10.1105/tpc.20.00099

Molin WT, Yaguchi A, Blenner M, Saski CA. Autonomous replication sequences from the Amaranthus palmeri eccDNA replicon enable replication in yeast. BMC Res Notes. 2020b:13:330. 10.1186/s13104-020-05169-0

Molin WT, Wright AA, VanGessel MJ, McCloskey WB, Jugulam M, Hoagland RE. Survey of the genomic landscape surrounding the 5-enolpyruvylshikimate-3-phosphate synthase (*EPSPS*) gene in glyphosate-resistant *Amaranthus palmeri* from geographically distant populations in the USA. Pest Manag Sci. 2018:74(5):1109–1117. 10.1002/ps.4659

Møller HD, Larsen CE, Parsons L, Hansen AJ, Regenberg B, Mourier T. Formation of extrachromosomal circular DNA from long terminal repeats of retrotransposons in *Saccharomyces cerevisiae*. G3: Genes, Genomes, Genet. 2016:6(2):453–462. 10.1534/g3.115.025858

Montgomery J, Morran S, MacGregor DR, McElroy JS, Neve P, Neto C, Vila-Aiub MM, Sandoval MV, Menéndez AI, Kreiner JM, et al. Current status of community resources and priorities for weed genomics research. Genome Biol. 2024:25:139. 10.1186/s13059-024-03274-y

Noguera MM, Porri A, Werle IS, Heiser JW, Brändle F, Lerchl J, Murphy B, Betz M, Gatzmann F, Penkert M, et al. (2022) Involvement of glutamine synthetase 2 (*GS2*) amplification and overexpression in *Amaranthus palmeri* resistance to glufosinate. Planta. 2022:256(3):57. 10.1007/s00425-022-03968-2

Norsworthy JK, Korres NE, Walsh MJ, Powles SB. Integrating herbicide programs with harvest weed seed control and other fall management practices for the control of glyphosate-resistant Palmer amaranth (*Amaranthus palmeri*). Weed Sci. 2016:64(3):540–550. 10.1614/WS-D-15-00210.1

Norsworthy JK, Ward S, Shaw D, Llewellyn R, Nichols R, Webster TM, Bradley K, Frisvold G, Powles S, Burgos B, et al. Reducing the risks of herbicide resistance: best management practices and recommendations. Weed Sci. 2012:60(SP1):31–62. 10.1614/WS-D-11-00155.1

Ou S, Su W, Liao Y, Chougule K, Agda JR, Hellinga AJ, Lugo CSB, Elliott TA, Ware D, Peterson T, et al. Benchmarking transposable element annotation methods for creation of a streamlined, comprehensive pipeline. Genome Biol. 2019:20:275. 10.1186/s13059-019-1905-y

Patterson EL, Saski CA, Sloan DB, Tranel PJ, Westra P, Gaines TA. The draft genome of *Kochia scoparia* and the mechanism of glyphosate resistance via transposon-mediated EPSPS tandem gene duplication. Genome Biol Evol. 2019:11(10):2927–2940. 10.1093/gbe/evz198

Picart-Picolo A, Grob S, Picault N, Franek M, Llauro C, Halter T, Maier TR, Jobet E, Descombin J, Zhang P, et al. Large tandem duplications affect gene expression, 3D organization, and plant–pathogen response. Genome Res. 2020:30(11):1583–1592. 10.1101/gr.261586.120

Powles SB, Yu Q. Evolution in action: plants resistant to herbicides. Annu Rev Plant Biol. 2010:61(1):317–347. 10.1146/annurev-arplant-042809-112119

Priess GL, Norsworthy JK, Godara N, Mauromoustakos A, Butts TR, Roberts TL, Barber T. Confirmation of glufosinate-resistant Palmer amaranth and response to other herbicides. Weed Technol. 2022:36(3):368–372. 10.1017/wet.2022.21

Raiyemo DA, Montgomery JS, Cutti L, Abdollahi F, Llaca V, Fengler K, Lopez AJ, Morran S, Saski CA, Nelson DR, et al. Chromosome-level assemblies of *Amaranthus palmeri*, *Amaranthus retroflexus*, and *Amaranthus hybridus* allow for genomic comparisons and identification of a sex-determining region. bioRxiv. 2024: 2024–09. 10.1101/2024.09.18.613719

Rice PA, Baker TA. Comparative architecture of transposase and integrase complexes. Nat Struct Mol Biol. 2001:8(4):302–307. 10.1038/86166

Robinson JT, Thorvaldsdóttir H, Winckler W, Guttman M, Lander ES, Getz G, Mesirov JP. Integrative genomics viewer. Nat Biotechnol. 2011:29(1):24–26. 10.1038/nbt.1754

Spier Camposano H, Molin WT, Saski CA. Sequence characterization of eccDNA content in glyphosate sensitive and resistant Palmer amaranth from geographically distant populations. PLoS One. 2022:17(9):e0260906. 10.1371/journal.pone.0260906

Steinrücken HC, Amrhein N. The herbicide glyphosate is a potent inhibitor of 5-enolpyruvylshikimic acid-3-phosphate synthase. Biochem Biophys Res Commun. 1980:94(4):1207–1212. 10.1016/0006-291X(80)90547-1

Sulzback JRM, Borgato EA, Cutti L, Hill EC, Burns EE, Patterson EL. Optimizing molecular assays for glyphosate and ALS-Inhibitor resistance diagnostics in four weedy species. Can J Plant Sci. 10.1139/cjps-2024-0020

Takano HK, Beffa R, Preston C, Westra P, Dayan FE. A novel insight into the mode of action of glufosinate: how reactive oxygen species are formed. Photosynth Res. 2020:144(3):361–372. 10.1007/s11120-020-00749-4

Turner KM, Deshpande V, Beyter D, Koga T, Rusert J, Lee C, Li B, Arden K, Ren B, Nathanson DA, et al. Extrachromosomal oncogene amplification drives tumour evolution and genetic heterogeneity. Nat. 2017:543:122–125. 10.1038/nature21356

Varanasi VK, Brabham C, Norsworthy JK. Confirmation and characterization of non-target site resistance to fomesafen in Palmer amaranth (*Amaranthus palmeri*). Weed Sci. 2018:66(6):702–709. 10.1017/wsc.2018.60

Wang K, Tian H, Wang L, Wang L, Tan Y, Zhang Z, Sun K, Yin M, Wei Q, Guo B, et al. Deciphering extrachromosomal circular DNA in Arabidopsis. Comput Struct Biotechnol J. 2021:19:1176–1183. 10.1016/j.csbj.2021.01.043

Zhang C, Johnson NA, Hall N, Tian X, Yu Q, Patterson EL. Subtelomeric *5-enolpyruvylshikimate-3-phosphate synthase* copy number variation confers glyphosate resistance in *Eleusine indica*. Nat Commun. 2023:14(1):4865. 10.1038/s41467-023-40407-6

Zhang C, Yu Q, Han H, Yu C, Nyporko A, Tian X, Beckie H, Powles S. A naturally evolved mutation (Ser59Gly) in glutamine synthetase confers glufosinate resistance in plants. J Exp Bot. 2022:73(7):2251–2262. 10.1093/jxb/erac008

Zhuang J, Zhang Y, Zhou C, Fan D, Huang T, Feng Q, Lu Y, Zhao Y, Zhao Q, Han B, et al. Dynamics of extrachromosomal circular DNA in rice. Nat Commun. 2024:15(1):2413. 10.1038/s41467-024-46691-0

Zuo S, Yi Y, Wang C, Li X, Zhou M, Peng Q, Zhou J, Yang Y, He Q. Extrachromosomal circular DNA (eccDNA): from chaos to function. Front Cell Dev Biol. 2022:9:792555. 10.3389/fcell.2021.792555

